# Unveiling inter individual variability of human fibroblast innate immune response using robust cell-based protocols

**DOI:** 10.1101/2020.06.04.133264

**Authors:** A. Chansard, N. Dubrulle, M. Poujol de Mollens, P. B. Falanga, T. Stephen, M. Hasan, G. van Zandbergen, N. Aulner, S. L. Shorte, B. David-Watine, for the *Milieu Intérieur* Consortium

## Abstract

The *LabEx Milieu Interieur* (*MI*) project is a clinical study centered on the detailed characterization of the baseline and induced immune responses in blood samples from 1000 healthy donors. Analyses of these samples has lay ground for seminal studies on the genetic and environmental determinants of immunologic variance in a healthy cohort population. In the current study we developed *in vitro* methods enabling standardized quantification of MI-cohort-derived primary fibroblasts responses. Our results show that *in vitro* human donor cohort fibroblast responses to stimulation by different MAMPs analogs allows to characterize individual donor immune-phenotype variability. The results provide proof-of-concept foundation to a new experimental framework for such studies.

A bio-bank of primary fibroblast lines was generated from 323 out of 1,000 healthy individuals selected from the MI-study cohort. To study inter-donor variability of innate immune response in primary human dermal fibroblasts we chose to measure the TLR3 and TLR4 response pathways, both receptors being expressed and previously studied in fibroblasts. We established high-throughput automation compatible methods for standardized primary fibroblast cell activation, using purified MAMPS analogs, poly I:C and LPS that stimulate TLR3 and TLR4 pathways respectively. These results were in turn compared with a stimulation method using infection by HSV-1 virus. Our “Add-only” protocol minimizes high-throughput automation system variability facilitating whole process automation from cell plating through stimulation to recovery of cell supernatants, and fluorescent labeling. Images were acquired automatically by high-throughput acquisition on an automated high-content imaging microscope. Under these methodological conditions standardized image acquisition provided for quantification of cellular responses allowing biological variability to be measured with low system noise and high biological signal fidelity. Optimal for automated analysis of immuno-phenotype of primary human cell responses our method and experimental framework as reported here is highly compatible to high-throughput screening protocols like those necessary for chemo-genomic screening. In context of primary fibroblasts derived from donors enrolled to the MI-clinical-study our results open the way to assert the utility of studying immune-phenotype characteristics relevant to a human clinical cohort.

## INTRODUCTION

The *LabEx Milieu Interieur* (*MI*) project (www.milieuinterieur.com) is a clinical study aiming to provide the first description of both genetic and environmental determinants of immunologic variance within the general healthy population (Thomas *et al.*, 2015). Central to this study, blood samples from healthy volunteers were stimulated by a range of 40 distinct stimuli allowing to characterize the induced immune response (*i.e.* major secreted cytokines chemokines, transcriptomr *etc. see* Duffy *et al.*, 2014; Urrutia *et al.*, 2016; Piasecka *et al.*, 2018). As part of the MI-program skin biopsies were collected from 323 out of 1,000 healthy individuals and primary human fibroblasts were prepared (Genethon, Evry, France). The MI-fibroblasts collection (323 primary fibroblasts) comprises a one-of-a-kind cell collection inasmuch as each derived primary cell isolate is uniquely associated with its corresponding donor data in the *LabEx MI* genotype-to-phenotype database (e.g. serology, genomic/proteomic analysis, microbiota, clinical data, and epidemiological selection criteria).

Fibroblasts are the principal cellular constituents of connective tissues. Functionally fibroblasts can be described as a population of cells that synthesize and secrete a complex array of structural and non-structural extra-cellular matrix (ECM) molecules. Fibroblasts actively organize and remodel ECM through the production of proteinases, and converse with nearby cells through paracrine, autocrine and other forms of communication (Sorrell *et* Caplan, 2009). As such fibroblasts are fundamental to tissue homeostasis and normal wound repair. The role of fibroblasts in the defense against pathogens and more generally in innate immunity has only more recently emerged with the discovery of Toll-like receptors (TLRs; Tabeta *et al.*, 2000). The TLRs family consists of more than 13 prototype pattern-recognition receptors (PRRs) that recognize microbial-associated molecular patterns (MAMPS) from various microbial pathogens such as viruses, bacteria, protozoa and fungi or from danger-associated molecular patterns (DAMPS) from damaged tissue (Kawai *et* Akira, 2007; Schaefer, 2014). Moreover, dominant producers of interleukin-6 (IL-6) at sites of peripheral inflammation, fibroblasts are now recognized for their major contribution to inflammatory regulation (Noss *et al.*, 2015; Nguyen *et al.*, 2017). Fibroblasts are emerging with a key role in local innate immunity evoked during response to pathogens as well as vaccines (Jolly *et al.*, 2017; Kübacher *et al.*, 2017; Salyer *et* David, 2018; LeBleu *et* Neilson, 2019).

The *LabEx* MI program highlighted the value of developing and systematically using standardized methodologies for measuring blood host immune responses. The systematic standardization of blood sample preparation and handling protocols enabled an unprecedented measure of inter-individual phenotype variance relevant to the human population cohort. Particularly, MI-studies showed that the whole blood stimulation by different MAMPs analogs are more informative to reveal the individual variability than unstimulated samples (Duffy *et al.*, 2014; Kim-Hellmuth *et al.*, 2017).

Our MI-program experience raised the fundamental question of wether the MI-fibroblast collection might also present innate immune response characteristics relevant to studying inter-individual donor variance.The first step to address this question requires to validate methods and protocols allowing to characterize and quantify the inter-individual variance of the innate immune response of the donor primary fibroblast collection.

That TLR3 and TLR4 receptors are integral to innate immune response in fibroblasts is supported by numerous studies (Wang *et al.*, 2011; Yao *et al.*, 2015). Further, deficits in the TLR3 pathway are associated with cases of herpes viral encephalitis in young children and can be identified *in vitro assay* with patient’s fibroblasts (Sancho-Shimizu *et al.*, 2011; Guo *et al.*, 2011; Zhang *et al.*, 2007). We therefore hypothetized that TLR3/TLR4 variability in the MI-population cohort should be detectable *in vitro* and might therefore provide an insightful measure of inter-individual variance preserved in fibroblasts of the MI-study. In a first step toward investigating such a phenomena in a high-value asset primary human cell collection comprising over three-hundred unique donor primary fibroblasts we report here the development of two standardized assays:

1. HSV-1(Herpes Simplex Virus type 1) infection: monitoring of the cell response were developed and tested by using cells harboring mutations in the TLR3 pathway. HSV-1 is an ubiquitous human neurotropic virus that affects up to 85% of the world’s population with recurrent clinical HSV-1 infections and manifestations (De Chiara *et al.*, 2012; Looker *et al.*, 2015). HSV-1 is recognized by different TLRs: TLR2 for surface viral structures, TLR3 for dsRNA, and TLR9 for viral DNA (Melchjorsen, 2012).
2. MAMPS analogs stimulation: the TLR3 ligand of choice is polyinosinic-polycytidilic acid (poly I:C), a synthetic analog of double-stranded RNA (dsRNA), a MAMP associated with viral infection. On the otherhand TLR4 is stimulated by lipopolysaccharide (LPS), a MAMP associated with the major structural component of the outer wall of all Gram-negative bacteria.

After recognition of MAMPs, TLRs recruit a cascade of adaptor proteins by homophilic protein-protein interactions *via* their TIR-domains (Toll/interleukin-1 receptor/resistance protein domain) leading to the activation of transcription factor nuclear factor-kappa B (NF-κB). Accordingly, both TLR3 and TLR4 pathways lead to translocation from the cytoplasm to the nucleus of the transcription factor NF-κB, albeit with different kinetics and *via* distinct protein-protein signaling cascades. The early phase of NF-κB activation depends on the MyD88-dependent pathway and the late phase NF-κB activation is controlled by the TRIF-dependent pathway (Kawai *et* Akira, 2007). We therefore reasoned visual detection of NF-κB nuclear translocation can be used to quantify TLR3/ TLR4 activation. Accordingly, we developed an “Add-only” protocol to avoid signal variability resulting from medium exchange. We validate the protocols for poly I:C and LPS stimulation by comparing responses in a subset of MI-donor primary fibroblasts (13 of the 323), to measurements performed on MI-extraneous immortalized fibroblasts lines from patients with mutations in the TLR3 or TLR4 pathways, and commercially supplied primary fibroblasts from single donors.

Hereby we describe our highly standardized protocols for stimulation/infection of primary human fibroblasts and the measurement of their immune response signatures at cellular and protein level which has enabled the identification of high and low fibroblast-specific responses to LPS.

## MATERIALS AND METHODS

### Cells and Donors

DHF primary fibroblasts (women, 55 years old, causasian, Zenbio Inc.) were purchased at Tebu, France. BJ (CRL-2522) and WI-38 (CCL-75) human fibroblasts were from ATCC.

The following SV40-immortalized fibroblasts were provided by J-L. Casanova, Laboratoire de génétique humaine des maladies infectieuses, Université Paris Descartes: TLR3^−/−^ (Boisson *et al.*, 2012), 2018 (TLR3^+/−^) and 1323 (TLR3^+/−^) (Zhang *et al.*, 2007), TRIF^−/−^(Sancho-Shimizu *et al.*, 2011), TBK1^+/−^(Herman *et al.*, 2012), Nemo^−/−^ (Smahi *et al.*, 2000) from patients and two control fibroblasts from healthy donors, C65 and C72.

*LabEx MI* primary fibroblasts were prepared from biopsies of the non-sun exposed interior of the arm of healthy volunteers at the Genethon (Evry, France). The identity of the subjects is coded by a number. The following numbers: 209, 220, 221, 241, correspond to women age 30-39; 318, 323, 341, 376 to men age 30-39; 818, 819, 820 to women age 60-69; and 914, 915: men age 60-69. Biopsies were obtained from a subset of the *LabEx* MI healthy donor cohort, approved by the *Comité de protection des personnes* – Ouest 6 (Committee for the protection of persons) on June 13th, 2012 and by the French Agence nationale de sécurité du médicament (ASNM) on June 22nd, 2012. The study is sponsored by Institut Pasteur (Pasteur ID-RCB Number: 2012-A00238-35) and was conducted as a single centre interventional study without an investigational product. The original protocol was registered under ClinicalTrials.gov (study# NCT01699893). The samples and data used in this study were formally established as the Milieu Interieur biocollection (NCT03905993), with approvals by the Comité de Protection des Personnes – Sud Méditerranée and the Commission nationale de l’informatique et des libertés (CNIL) on April 11, 2018.

### Cell culture

The cells were cultured in DMEM GlutaMAX (Dulbecco’s Modified Eagle Medium, Life Technology) supplemented with 10% FCS (fetal calf serum, Life Technology) without antibiotics. The cells were incubated at 37°C, 5% CO_2_. Primary fibroblasts were passaged at 70-80% confluency. The cells are said to pass n + 1 each time they are detached by trypsinization and seeded again in new culture flask with fresh medium. For primary dermal human fibroblasts, one passage corresponds in average to the multiplication by three of the cell population over a period of one week. Routinely, the cells were analyzed between passage 5-6 and passage 10.

Cells were regularly assessed for mycoplasma contamination by using the Mycotest kit (Enzo).

### Reagents

For infection experiments, we used a HSV-1 in which GFP (green fluorescent protein) was fused to a viral capsid protein (VP26) (strain KOS, Desai *et* Person, 1998). A high concentration stock of HSV-1-GFP was a generous gift from Prof. Desai (Virology Laboratories, Department of Pharmacology and Molecular Sciences, Johns Hopkins University School of Medicine, Baltimore, Maryland).

Poly I:C (High molecular weight), a synthetic analogue of double-stranded RNAs that binds to the TLR3 receptor, and LPS (LPS-EB Ultrapure E.coli 0111: BA), a component of the gram-negative bacterial membrane that binds to the TLR4 receptor were purchased from Invivogen.

Human TNF*α* premium grade was from Miltenyi Biotec and Interferon *α*-2b INTRONA from Merck MSD France.

### Immuno-fluorescent labeling

To carry out the immuno-fluorescent labeling, the cells were permeabilized for 15 minutes in PBS-0.1% Triton (D-PBS, Life Technology). The cells were then incubated for 1 hour at room temperature in blocking buffer (PBS-fat dry milk 5%). Then, the cells were incubated with a monoclonal anti-human NF-B p65 antibody (27F9.G4, 1/2000, Rockland) diluted in the blocking solution (PBS-milk 5%) overnight at 4 °C. Fluorescent anti-mouse secondary antibodies were diluted in the blocking solution and incubated for 2 hours at room temperature. Finally, the nuclei were labeled with Hoechst 33342 (0.5 μg / ml, Invitrogen) for 15 minutes at room temperature.

### Infection with Herpes Simplex Virus-1

At day 1, the cells were plated in a 96-well plate (μClear Bottom, Greiner CAT 655090): 12,000 cells per well for primary fibroblasts and 6,000 cells per well for fibroblast lines immortalized in 50 μl of cell culture medium. The cells were incubated for 24 hours before being infected at different multiplicities of virus (MOI) (0.01, 0.05, 0.1 or 0.5 MOI) in DMEM at 2% FCS. The virus was left in contact with the cells for two hours, then was replaced by complete DMEM medium at 10% FCS. The cells were then incubated for a further 24 hours at 37 °C and 5% CO2. The cells were then fixed in 4% PFA (Electron Microscopy Sciences) for 15 minutes at room temperature. Finally, the nuclei of the cells were labeled with Hoechst 33342. The GFP fluorescence of the samples were quantified at 24 hours after infection. For assaying cell protection to viral infection, cells were treated with 105 IU/ml IFN *α*-2b for 18 hours before infection.

### Protocol “Add-only” for poly I:C and LPS stimulation

At day 0: The cells were plated in 100 μL of DMEM 10% FCS in a 96-well plate (Clear Bottom, Greiner): 12,000 cells per well for primary fibroblasts and fibroblast lines and 6,000 cells per well for the immortalized fibroblast lines.

At day 1: Two times concentrated poly I:C : 0.2 μg, 2 μg, 10 μg and 20 μg for 1 ml; or two times concentrated LPS: 0.2 μg, 1 μg, 2 μg of LPS for 1 ml, were prepared in DMEM 10% FCS and 100 μL were added on top of the cells without medium withdrawing.

Incubation time was 3 hours for LPS and 22 hours for poly I:C for monitoring the expression of NF-κB.

At day 2: Twenty-five μl of PFA 32% are added to the 200 μl of medium to achieve a final concentration of 3,5%. After 15 min, the cells were carefully rinsed with PBS twice then processed for immunofluorescent labelling.

For cytokine analysis, the supernatants were retrieved 24 hours after the beginning of the stimulation and transferred to a plate for freezing.

All the steps of this protocol can be easily adapted to automation (data not shown).

### Image acquisition and analysis

After immunofluorescent labelling of fixed cells, two channel images were acquired in a fully automated and unbiased manner using an automated spinning disk confocal microscope (OPERA QEHS, Perkin Elmer Technologies, UtechS PBI, Institut Pasteur) and a 10× air objective (NA=0.4) with the following sequential acquisition settings: (i) 561 nm laser line excitation, filter 600/40 for *Cy3* detection, or 488 nm laser line excitation, filter 540/75 for Alexa 488 detection and (ii) 405 nm laser line excitation, filter 450/50 for Hoechst 33342 detection. Forty-seven images per channel, covering roughly the entire surface of each well, were collected for reliable statistical analysis considering potential spatial cell and compound distribution biases.

The images and associated metadata were transferred to the Columbus Conductor™ Database (Perkin Elmer Technologies) for storage and further analysis. The image analysis was performed by batches in Columbus using custom designed image analysis building blocks.

For HSV-1 infection, only the fluorescence intensity in nuclei containing a green fluorescent protein (VP26-GFP) fused capsid protein is considered to identify HSV-1 positive cells. A ratio of cells defined as HSV-1 (GFP) positive is calculated on the total population defined by the number of Hoechst labeled nuclei, which corresponds to the percentage of infected cells on the total population at a certain multiplicity of infection.

To analyze the translocation of NF-κB in the nucleus, the routine allows to enumerate the total number of cells and to measure the average intensities of fluorescence *per* pixel (or unit area) in the masks considered (the nucleus and the cytoplasm) in each individual cell. A ratio of the fluorescence intensity of the nucleus to the fluorescence intensity of the cytoplasm is calculated: if this ratio is higher at 1.2 we consider that the cells are stimulated, if this ratio is less than 1 we consider that the cells have not been stimulated. We report then the proportion of stimulated cells in the entire population of each well at a given condition to characterize the stimulus level.

### Cytokine flow cytometry

Cells were stimulated with LPS 1μg/ml or poly I:C 20μg/ml during 72hours, then treated with brefeldin A 1x (Abcam ab193369) for 6 hours. The cells were trysinized and resuspended in 0,5 ml PBS and fixed by addition of 0,5 ml PBS 8% PAF on ice for 20 min. Fibroblasts were then permeabilized in PBS-SVF1%, 0,3% Triton X100 for 20 min at 4°C. The cells were labelled in PBS-SVF1%, 0,1% Triton X100 with anti-huIL-6-FITC (BD Biosciences, 340526) and anti-huIL-8-APC (Miltenyi Biotec, 130-112-015) and analyzed on a Fortessa cytometer in the Cytometry and Biomarkers UtechS, Institut Pasteur.

### Cytokine dosage

Cell culture media were collected post-stimulation at different timepoints and stored at −80°C until further analysis. The thawed samples were centrifuged at 13 000g at 4°C for 10 min. In the undiluted media samples, the cytokine levels were determined by Luminex xMAP technology using Human Custom ProcartaPlex Assay kit (Thermofisher, Cat PPX-04-MXRWEYU) in accordance with the manufacturer’s recommendations. The protocol was adapted to DropArray plate (Clinisciences, Cat 96-CC-BD-05); a luminex method based on the use of a 96 wall-less plate capable of a five-fold miniaturized format with regards to sample (5 μL minimum) and Luminex reagents (Le Guezennec *et al.*, 2015). The cytokine levels were measured for a combination of 4 different cytokines : interleukin-6 (IL-6), interleukin-8 (IL-8), interferon α (IFN α) and Monocyte Chemotactic Protein-1 (MCP-1). Plate was read on the BioPlex 200 (BioRad). Concentration were extrapolated with the Bioplex Manager software (v6.1) using 5-PL curve-fitting regression algorithms with standards run in duplicate. For data representation, out of ratio values in high concentration (OOR >) were replaced with a value that is twice of the upper limit of quantification (ULOQ), which is measured in the dataset based on the standard curve. Cytokine dosage was performed at the Cytometry and Biomarker UtechS, Institut Pasteur.

### Statistical analysis

Each experiment was repeated at least twice with every experimental point aggregated from triplicate sample measurement.

Mean values and standard deviation were calculated for all results.

To compare LPS high and low responders, the arithmetic means were calculated for each examined group. Comparisons between the two groups were analyzed by Mann–Whitney-U-Test. Probability values (p) were calculated and a level of *P* < 0.05 was considered as statistically significant.

Chi-square test was used for univariate analysis of percentage of cells with NF-kB in the nucleus of the cells. Data were presented as means ± standard deviation from at least three independent experiments. Statistical significance was defined as *P* < 0.05.

### Consortium

The *Milieu Intérieur* Consortium is composed of the team leaders: Laurent Abel (Hôpital Necker), Andres Alcover, Hugues Aschard, Kalla Astrom (Lund University), Philippe Bousso, Pierre Bruhns, Ana Cumano, Caroline Demangel, Ludovic Deriano, James Di Santo, Françoise Dromer, Gérard Eberl, Jost Enninga, Jacques Fellay (EPFL, Lausanne), Ivo Gomperts-Boneca, Milena Hasan, Serge Hercberg (Université Paris 13), Olivier Lantz (Institut Curie), Hugo Mouquet, Etienne Patin, Sandra Pellegrini, Stanislas Pol (Hôpital Côchin), Antonio Rausell (INSERM UMR 1163 – Institut Imagine), Lars Rogge, Anavaj Sakuntabhai, Olivier Schwartz, Benno Schwikowski, Spencer Shorte, Frédéric Tangy, Antoine Toubert (Hôpital Saint-Louis), Mathilde Trouvier (Université Paris 13), Marie-Noëlle Ungeheuer, Darragh Duffy§, Matthew L. Albert (In Sitro)§, Lluis Quintana-Murci§,

¶ unless otherwise indicated, partners are located at Institut Pasteur, Paris

§ co-coordinators of the Milieu Intérieur Consortium.

Additional information can be found at http://www.milieuinterieur.fr/

## RESULTS

### 1. Analysis of HSV-1 cell infection

As a means to evaluate the variability of the innate immune response to infection of fibroblasts from different donors, we first developed an assay using HSV-1 infection using virus engineered to express green fluorescent protein (GFP). The GFP from *Aequorea victoria* was fused with the capsid protein VP26 to generate a VP26-GFP protein expressed by the K26GFP virus. VP26 is located on the outer surface of the capsid and cells infected with the K26GFP virus exhibited a punctate nuclear fluorescence at early times in the replication cycle. At later times during infection a generalized cytoplasmic and nuclear fluorescence, including fluorescence at the cell membranes and the spreading of infection to surrounding cells was observed (Desai *et* Person, 1998). To our knowledge, the combination of imaging of a fluorescent signal in a high content setting to follow HSV-1 infection efficiency has not yet been described.

To evaluate the delay between HSV-1 infection and the appearance of the VP26-GFP protein in infected cells, we first performed automated imaging microscopy using the IncuCyte Zoom device (Essen Biosciences). The fluorescent signal corresponding to the production of the capsid protein-GFP (VP26), produced after a replication cycle (Desai and Person, 1998), appeared only about 9-10 hours after infection in our experimental conditions. This approach allowed to observe viral replication dynamics by following the fluorescent signal which appearing more rapidly indicated increased susceptibility of cells to infection (data not shown).

Our aim was to establish a robust end-point measure that would allow a quantitative assessment of the response to infection. We thus tried a large range of multiplicity of infection (from 0,004 to 10 MOI) and several incubation times after infection to determine the conditions where GFP expression was optimally detected. Based on the range of the number of infected cells detected, we chose to image the infection at 48 hours post infection at between 0,01 to 0,5 MOI (data not shown).

We first followed the infection of two different cell lines (DHF and TLR3−/−). According to published data, our hypothesis in developing these protocols was that DHF cells, considered to be healthy fibroblasts, should be able to respond efficiently to HSV-1 infection due to the stimulation of the TLR3 receptor and therefore be less infected than the the TLR3_−/−_ fibroblast cell line which are deficient in their TLR3 pathway, leading to increased sensitivity to infection. As shown in figure 1A, the infection process can be followed through expression of GFP in the nucleus of the infected cells. Active infection of DHF cells could be detected at 48 hours for an MOI of 0,01 or higher (data not shown). Image analysis performed as described in Materials and Methods, showed that at 0,1 MOI, 80-90% of DHF cells are infected versus only 40-50% for TLR3_−/−_ cells. At 0,5 MOI 90 to 95% of DHF and TLR3_−/−_ cells are infected (Figure 1B). Thus, unexpectedly TLR3−/− cells appeared to be less sensitive than DHF cells at a 0,1 MOI infection by HSV-1 and to have comparable sensitivity at 0,5 MOI. By way of positive control, we also verified that treatment with interferon α-2b (10_5_ UI/ml) prior to infection totally protected the cells (data not shown).

**Figure 1:**
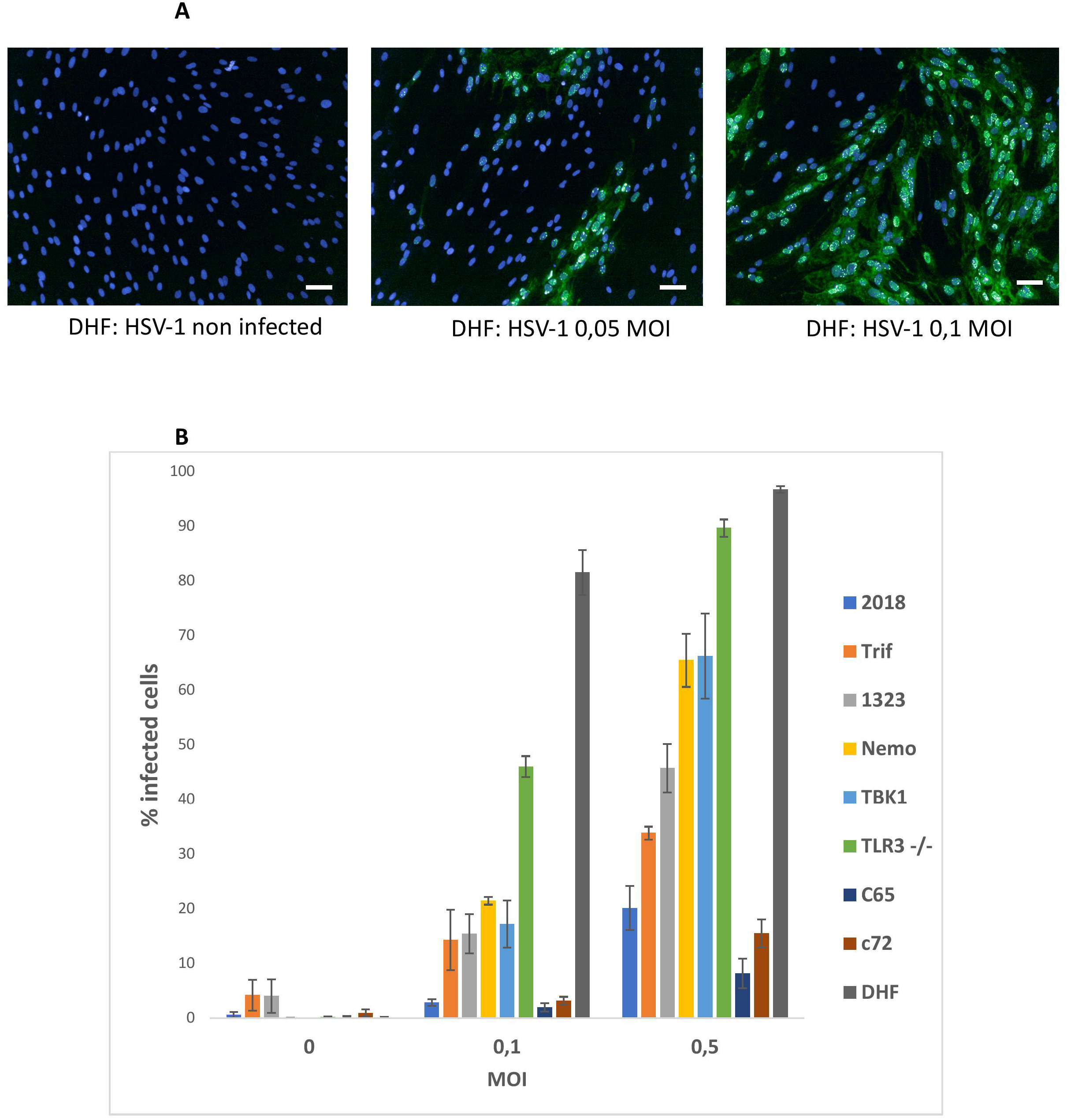
HSV1-GFP infection of healthy and mutant fibroblasts. A : Overlays of the detection of the fluorescence signal in DHF cells (green) infected at 05, 0,1 MOI and non-infected controls with a nuclei Hoechst counterstain (blue). Images were acquired on an Opera QEHS system with a 10x objective and correspond to one field of view. Scale bar: 50μm. B : Quantification of the percentage of infected cells after infection with 0, 01 and 0,5 MOIs. Compiled results of the mean percentages presented together for comparison: 2018 (TLR3 +/− cells), Trif (TRIF−/− cells), 1323 (TLR3 +/− cells), Nemo (NEMO −/− cells), TBK1 (TBK1 −/− cells), TLR3−/− cells, control cells C65 et C72, and DHF. Bars plot the mean ± sd of triplicates of each experimental point.

We then analyzed a series of human immortalized patient fibroblasts harboring various mutations of the TLR3 pathway described by J-L. Casanova and collaborators and impacting the sensitivity to HSV-1 infection patients (Sancho-Shimizu *et al.*, 2011; Guo *et al.*, 2011; Zhang *et al.*, 2007). By contrast with DHF cells, C65 and C72 cells, which are immortalized healthy fibroblasts used as controls, appeared to have very low (about 10-15%) levels of infection even at 0.5 MOI (Figure 1B).

Mutant cells for TLR3 _+/−_ (1323), TRIF, TLR3 _+/−_ (2018), Nemo and TBK1 showed a higher susceptibility to infection with respectively 25%, 35%, 50%, and 60% of infected cells at 0.5 MOI (Figure 1B). Unexpectedly, immortalized human fibroblasts from healthy donors are less sensitive to HSV-1 infection compared to DHF, a primary fibroblast, which may indicate that immortalization would confer resistance to HSV-1 infection. Indeed, two other widely used non-immortalized healthy fibroblast cell lines, WI-38 and BJ, have a sensitivity to HSV-1 infection similar to that of TLR3 − / − cells (Figure S1).

### 2. Stimulation of cells by poly I:C

As for the *LabEx* MI whole blood study (Duffy *et al.*, 2014), we searched for stimulation conditions providing defined molecular stimuli of innate immune responses with viral or bacterial MAMPs. Based on published data (Audry *et al.*, 2011; Guo *et al.*, 2011) on poly I:C stimulation of fibroblasts, we tried the concentrations ranging from 10 to 100μg/ml for 22 hours incubation time using in parallel, primary and immortalized fibroblasts. Immortalized fibroblasts with mutations of the TLR3 pathway were also used in order to better evaluate the range of sensitivity for our read-out of cell activation, the percentage of cells having translocated NF-κB to the nucleus.

We next determined the conditions for optimal activation of the primary fibroblasts DHF by poly I:C in comparison with TLR3 _−/−_ cells. TLR3 _−/−_ cells displayed no response to poly I:C stimulation (10 μg/ml) whereas DHFs cells were activated (Figure 2A), 60-70% of the cells showing a nuclear NF-κB signal (Figure 2B). The immortalized C65 and C72 fibroblasts from healthy individuals were used as controls for the mutant cells. As shown in Figure 2B, the C65, C72 and DHFs cells were maximally activated (70-90 %) at a concentration of 5 μg /ml to 10 μg /ml poly I:C, the highest concentration evoking responses in around 80-90% of cells. Under the same conditions TLR3 mutant cell lines were generally less activated by poly I:C. The NF-κB protein was very little or not at all translocated in the nuclei of the cells carrying mutation in the Nemo or TRIF proteins (5-25 %) and the NF-κB translocation was only observed in 30-40% of the cells carrying TBK1 mutation. Consistent with their genotype TLR3 _−/−_ double mutant cells did not respond at all to poly I:C and 1323 and 2018 both heterozygous for TLR3 mutation (TLR3 _+/−_) showed a low response (15% and 40%, respectively; Figure 2B).

**Figure 2:**
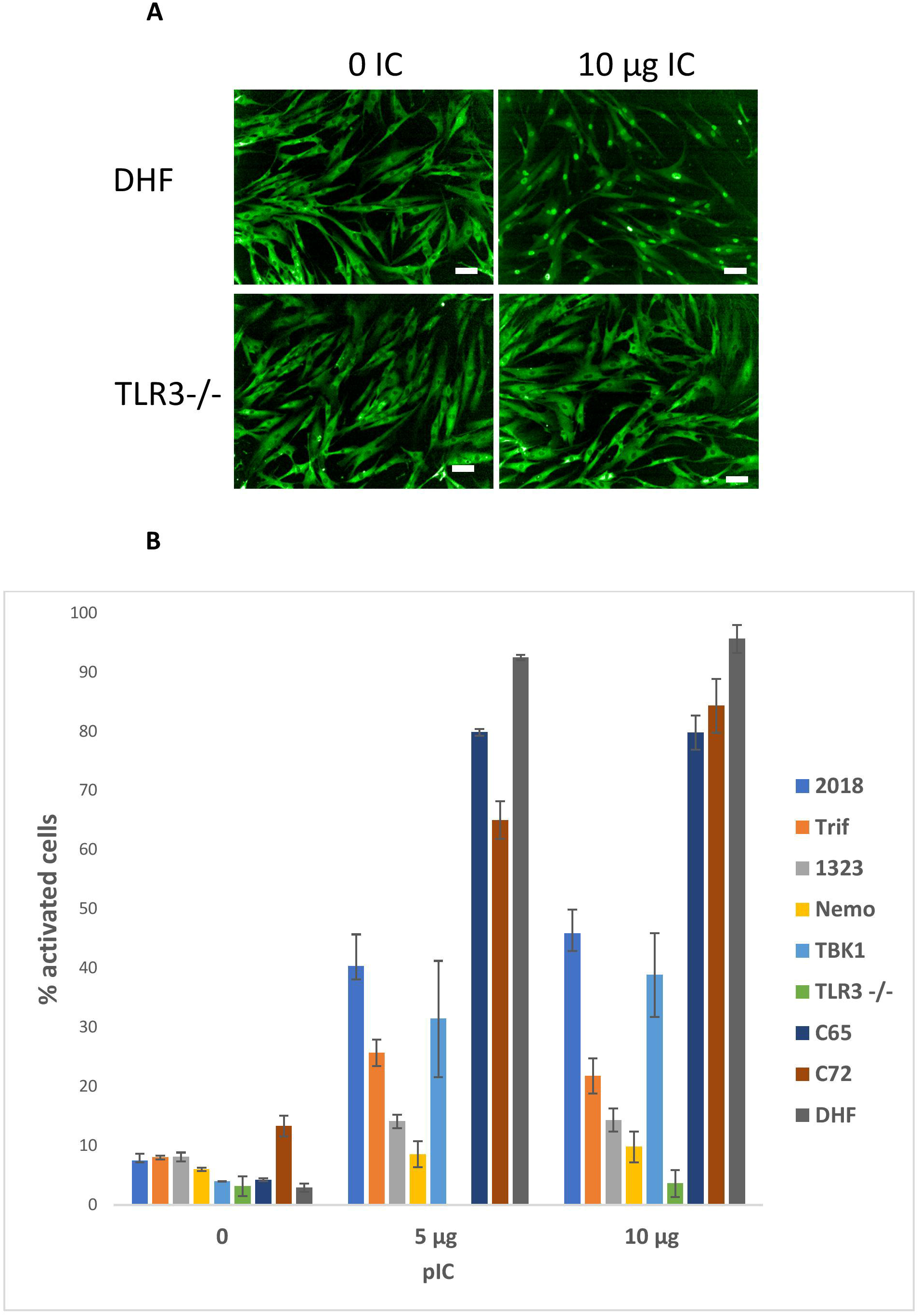
Poly (I:C) stimulation of healthy and mutant fibroblasts. A: Representative images of the immunofluorescence signal of DHF (upper part) and TLR3−/− (lower part) fibroblasts labelled with an anti-NF-κB antibody: non-stimulated cells (left) and stimulated with 10μg Poly (IC). Scale bar: 50μm. B: Percentage of cells with nuclear NF-κB (activated) for cells stimulated with 1μg and 10μg of Poly (I:C) and control unstimulated cells (0) : 2018 (TLR3 +/− cells), Trif (TRIF−/− cells), 1323 (TLR3 +/− cells), Nemo (NEMO −/− cells), TBK1 (TBK1 −/− cells), TLR3−/− cells, control cells c65 et c72, and DHF. Bars plot the mean ± sd of triplicates of each experimental point.

### 3. Variability of the response of LabEx MI primary fibroblasts to poly I:C and LPS

We then studied the response to poly I:C and LPS stimulation of a series of of thirteen primary fibroblasts from the *LabEx MI* using the “Add-only” protocol. As for poly I:C, we first evaluated the effect of different concentrations of LPS and incubation time on the activation of the cells defined as cells having translocated NF-κB from the cytoplasm to the nucleus. The incubation time was in the range of 1 to 3 hours, the highest signal following 3 hours incubation.

Using LPS concentrations ranging from 10 to 100 ng, we observed that the number of responding cells increased with the concentration and reached a maximum of 80-90% activated cells for some of the donors at a concentration of 100 ng/ml. Other cells were activated in the range of 30 to 50% and one donor fibroblast sample (323) showed no response at all. The figure 3A presents the combined mean and standard deviation measured from different experiments represented together for the comparison.

**Figure 3:**
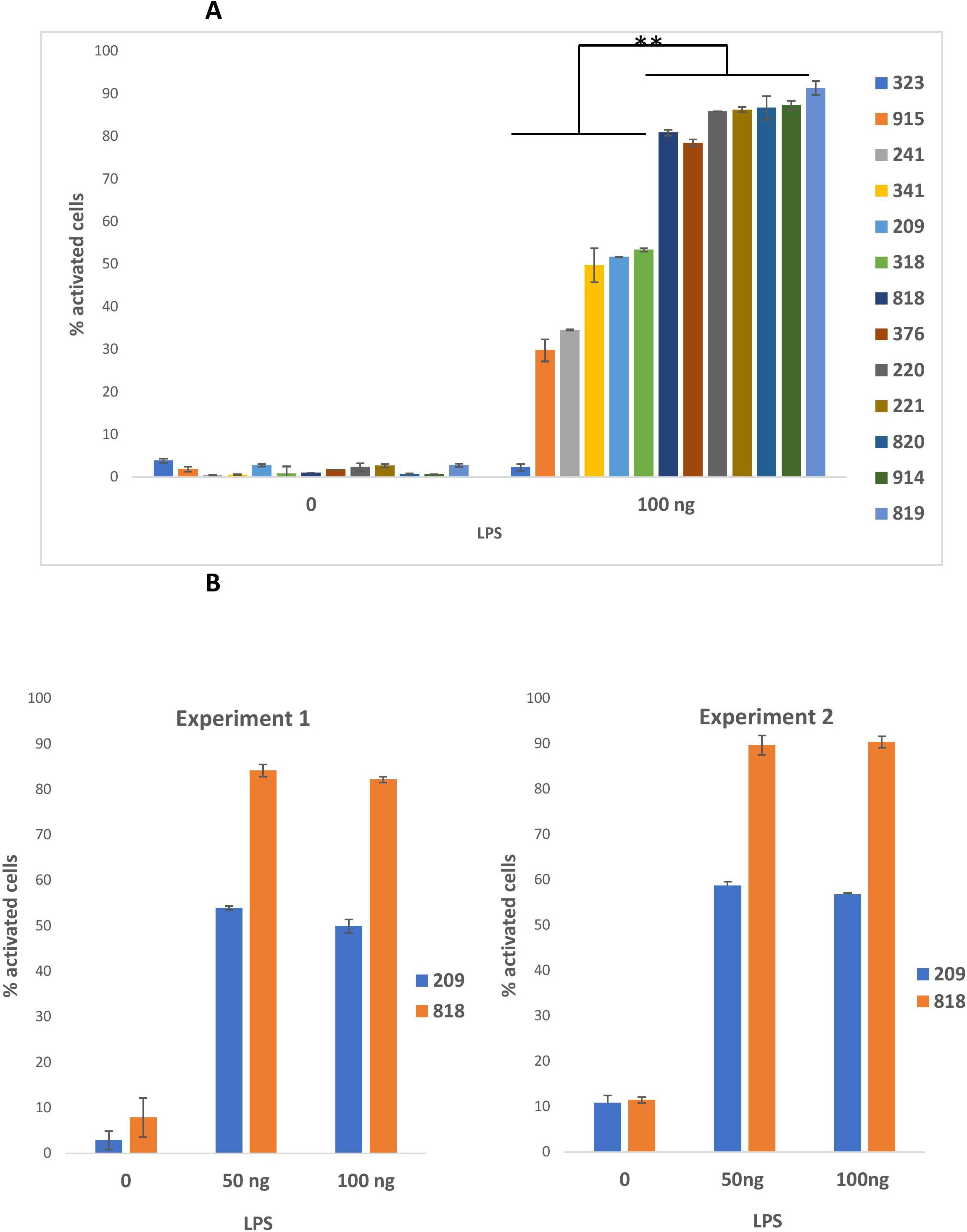
LPS stimulation of healthy fibroblasts. A:LPS and Poly (I:C) stimulation of 13 primary fibroblasts from the *LabEx Milieu interieur* collection: Percentage of cells with nuclear NF-κB (activated): compiled results of the mean percentages presented together for comparison for a stimulation with 100 ng of LPS and unstimulated controls. Bars plot the mean ± sd of triplicates of each experimental point. Statistical analyses were carried out using the Mann–Whitney-U-Test and p values < 0.05 were considered as statistically significant. ** Significant differences between low and high responders. B: Percentage of cells with nuclear NF-κB (activated) for donors’ fibroblasts (209, 818) activated with 50 and 100 ng of LPS or non-stimulated (0): results of two experiments performed at one year and a half apart. Bars plot the mean ± sd of triplicates of each experimental point. Chi-square test analysis of the results showed that differences between the 2 experiments were statistically not significant at a p value < 0,05.

We evaluated the robustness of the LPS cellular responses to be preserved characteristically by measuring the reproducibility over time of the cell response in two cell samples. The same experiment was performed with two donor samples (209 and 818) at an 18 month interval. In the first experiment, 209 and 818 cells were both at passage 8. In the second experiment, 209 cells were at passage 14 and 818 cells at passage 11 (Figure 3B). The relatedness of the two experimental results was evaluated by a *X*_i2_ test. There was no significant difference at a p-value superior to 0,05. Thus, the results were essentially the same.

We next assessed the response of the non-responder (323), a “low” responder (318) and two “high” responder (818 and 820) donor samples, as defined from the percentage of cells with nucleus NF-κB after LPS stimulation, to the stimulation by LPS (Figure 4A), poly I:C (Figure 4B) and TNF*α* (data not shown). All the cells were very similar in their responses to poly I:C and TNF*α* (data not shown), the response to poly I:C being a little less pronounced for 323 and 318 compared to the two other cells. Therefore, the deficient response of 323 cells appeared to be specific to the MyD88 arm of the TLR4 pathway.

**Figure 4:**
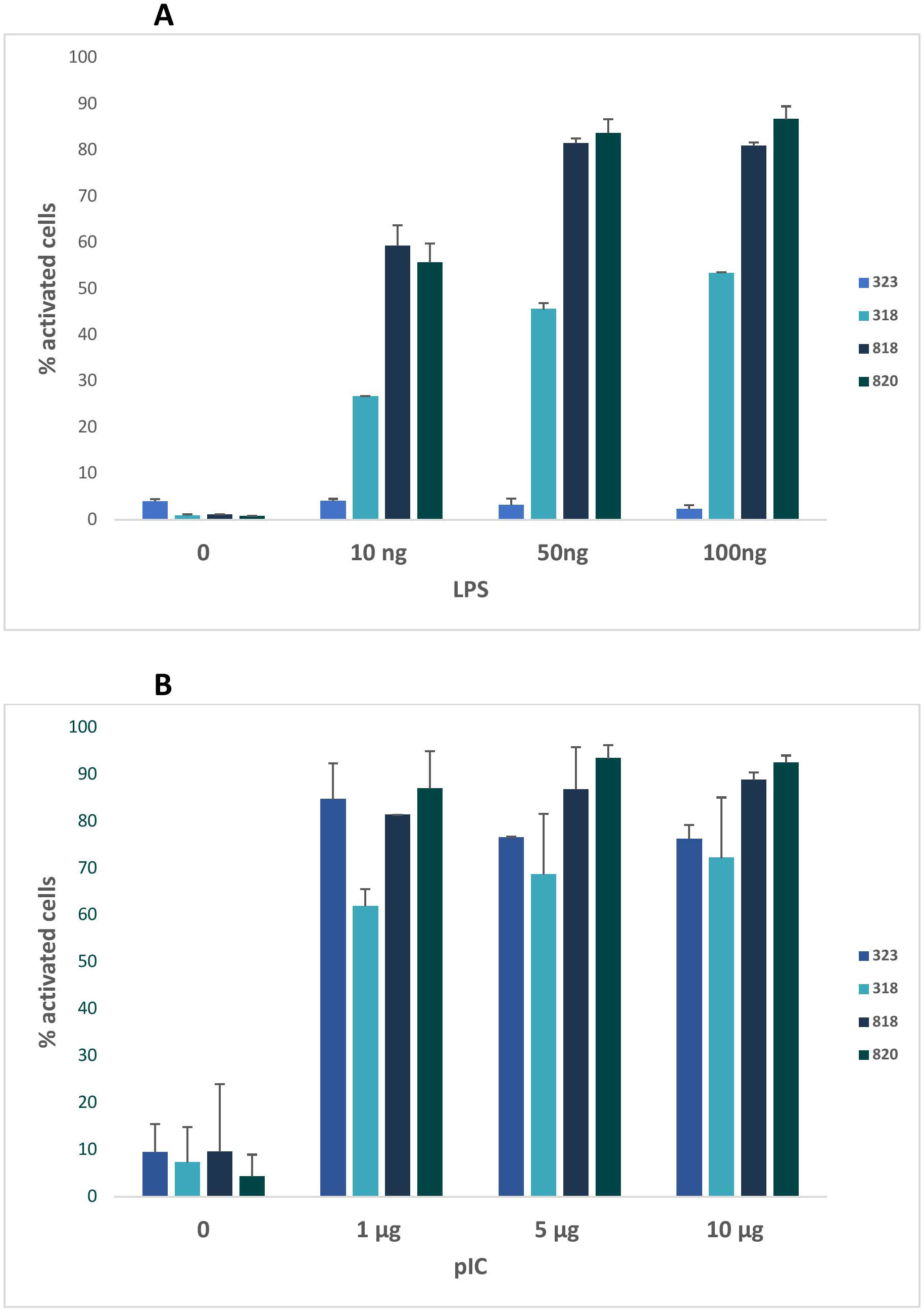
poly (I:C) and LPS responses of 323 cells compared to 318, 818 and 820 cells. A: Percentage of cells with nuclear NF-κB (activated) for donors’ fibroblasts (318, 323, 818, 820) activated with 10, 50 and 100 ng of LPS or non-stimulated (0). B: Percentage of cells with nuclear NF-κB (activated) for donors’ fibroblasts (318, 323, 818, 820) activated with 0,1, 1, 5 and 10 μg of Poly (IC) or non-stimulated (0). Bars plot the mean ± sd of triplicates of each experimental point.

We also evaluated the effect of the number of passages of primary fibroblasts on their response to stimulation. Indeed, it was generally accepted that fibroblasts must be analyzed before passage 10 at the latest and that it was preferable to compare the behavior of cells studied with equivalent passages.

We repeated the same experiment with cells at different passages. We observed that the response of the cells was not affected by the number of passages between 4 and 9 for poly I:C concentrations of 5 to 20 μg/ml (data not shown) and 8 to 14 as already shown for LPS stimulation (Figure 3B).

### 4. Flow cytometry and cytokine dosage of poly I: C and LPS stimulated fibroblasts

When comparing the response of the cells to various stimulations, we observed that the responses to LPS stimulation were distributed between “low” responder and “high” responders, while the inter-donor variability in the response to poly I:C was very low and the response to TNFα always involved 100% of the cells. To better evaluate the biological significance of the information consisting in the number of cells with NF-kB in the nucleus after stimulation, we stimulated fibroblasts from donor 209 (a LPS “low” responder), and donor 818, (a LPS “high” responder), with LPS and poly I:C measuring IL-6 and IL-8 in the cell culture supernatants. The expression of IL-6 was much strongly augmented in LPS stimulated cells from donor 818, but not cells from donor 209; whereas IL-8 was stimulated in both donors’ fibroblasts stimulated by LPS, but significantly more in cells from donor 818 than in donor 209. By contrast, stimulation by poly I:C induced a similar augmentation of both cytokines in the cell culture supernatants from both donors 209 and 818 (Figure 5 A).

**Figure 5:**
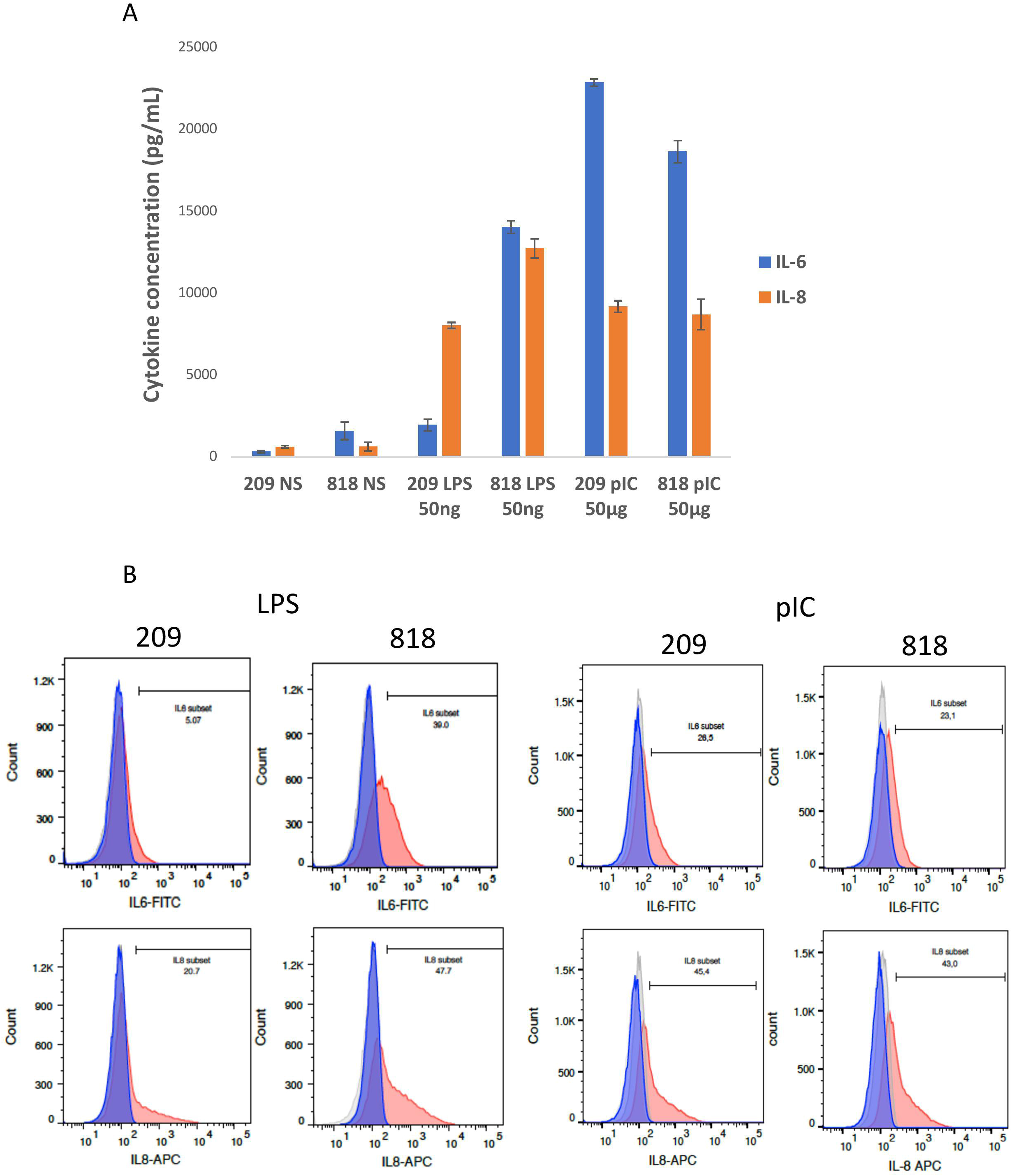
Analysis of cytokine expression by FACS and Luminex dosage. A: Flow cytometry analysis of IL-6 and IL-8 expressed by fibroblasts after LPS or poly (I:C) stimulation and control non-stimulated cells: the light grey histogram corresponds to non-labelled non-stimulated cells, the dark grey to stimulated non-labelled cells, the blue histogram represents non stimulated labelled cells and the red histogram cells stimulated and labelled. The bar indicates the percentage of labelled cells in stimulated conditions compared to controls. On the left: comparison of 209 and 818 cells stimulated by LPS; on the right: comparison of 209 and 818 cells stimulated by poly (I:C). B : IL-6 and IL-8 cytokine dosage in the supernatant of 209 and 818 stimulated by LPS and poly (I:C) and non-stimulated controls. Bars plot the mean ± sd of triplicates of each experimental point. Concentrations are given in pg/ml.

In parallel we analyzed the cellular expression levels of IL-6 and IL-8 in LPS and poly I:C stimulated samples prepared from donors 209 and 818 using single cell flow cytometry analysis. We observed that there were fewer cells expressing IL-6 and IL-8 cytokines in the LPS stimulated LPS “low responder” fibroblasts (donor 209) than in the LPS “high responder” (donor 818) (5% versus 39% for IL-6 and 20.7% versus 47,7% for IL-8, figure 4B, left). By contrast, there was little difference between the responses of the two populations of fibroblasts stimulated by poly I:C (26,5% versus 23,1% for IL-6 and 45,4% versus 43% for IL-8, figure 4B, right). Thus fibroblasts prepared from the donors 209 and 818 followed the same LPS and poly I:C response trend in both single cell cytokine expression, and cytokines released into the supernatant (figure 4).

## DISCUSSION

In this study, we report methods and protocol validating the proof-of-concept for quantitative cell-based assays useful to evaluate biologically relevant innate immune response in primary fibroblasts.

First, we used imaging of a fluorescent signal in a high content setting to follow HSV-1 infection efficiency. The assay is applicable to medium to high-throughput screening for anti-viral compounds as well as basic studies of cellular antiviral signaling. The observed responses were consistent with the data published for these mutants of the TLR3 pathway to HSV-1 infection (Sancho-Shimizu *et al.*, 2011; Audry *et al.*, 2011; Herman *et al.*, 2012; Guo *et al.*, 2011). Using this same protocol, we also observed that primary and immortalized human fibroblasts from healthy donors had different susceptibilities to HSV-1 infection. The fibroblasts used in our experiments were immortalized using the expression of the virus SV40 Large T (LT) antigen. SV40 LT interacts with endogenous proteins of the cells to induce immortalization, notably with the retinoblastoma protein, Rb, and with p53. In epithelial cells it was shown that Rb with E2F1 and p53 regulates the innate immune receptor TLR3 (Taura *et al.*, 2008; Taura *et al.*, 2012). In addition, immune response genes including interferon-stimulated genes are upregulated in cells expressing SV40 LT (Rathi *et al.*, 2010). This upregulated innate immune response may therefore explain the lower sensitivity of immortalized cells to HSV-1 infection, particularly at low MOI. These observations emphasize the importance of working with primary cells in place of SV40 LT immortalized cells to study the response to immune stimulants.

We developed an “add-only” protocol for stimulating cells by MAMPs and cytokines that minimized sample manipulation (no washing steps) that can give rise to well-to-well variability. This protocol was applied to three different cell types, primary fibroblasts, commercial lines and immortalized fibroblasts carrying mutations of the TLR3 pathway and allowed to differentiate individual responses based on the agonist concentration to which the cells respond. Inasmuch as cells displaying a low response level to poly I:C appeared to be inversely highly responsive to HSV-1 infection, both assays allowed to rank the innate immune response of these cells that involves the TLR3 pathway.

Of particular interest to us, we measured significant inter individual variability of the response to innate immune stimuli in primary fibroblasts from the MI-collection (percentage of activated cells with nuclear NF-κB) principally in response to LPS stimulation. Hundred per cent of the cells always responded to stimulation by TNFα and a low-range variability was observed in the response to poly I:C. By contrast, responses to LPS stimulation could be distributed in “low” and “high” responders. These different responses to poly I:C, LPS or TNFα probably reflect the efficiency of the pathways elicited by the corresponding receptors on NF-κB activation.

No differences related to age or gender were observed among the thirteen MI-cohort donors. High responses in the older donors could be a consequence of cell aging, a phenomenon called inflammaging (Freund *et al.*, 2010), however high responses are also observed in cells from younger donors. The observed responses are robust as we obtained essentially the same results for LPS stimulation of two cells (209 and 818) performed at an interval of eighteen months. Altogether, these results demonstrate that the “add-only” protocol actually allows to describe the inter individual variability of the innate immune response of human primary fibroblasts in culture. In addition, the cells of one donor (323) out of thirteen, a healthy donor with average range of blood responses to LPS, did not respond to LPS, but were stimulated by poly I:C and TNFα. We verified that this donor carried no SNPs in the TLR4 that were described to alter the response to LPS stimulation (Figueroa *et al.*, 2012; Long *et al.*, 2014). Therefore, these results are suggestive of a defect of the MyD88 arm of the TLR4 signaling in the donor 323 cells. It would be interesting to stimulate the 323 cells with other TLR agonists also using the MyD88 adaptor.

This is to our knowledge the first description of inter individual variability of the response of human primary fibroblast from healthy donors to innate immune stimulation. Individuals who consistently produced high or low concentrations of cytokines (most notably IL-1β, IL-6, IL-8 and TNF-α) in whole-blood samples after LPS stimulation had been previously identified. These phenotypes were stable, and corresponded to significant gene expression differences (Wurfel *et al.*, 2005).

Similarly to our results, analysis of the variability of cattle dermal fibroblasts responses to LPS allowed *low responder* (LR) and *high responder* (HR) classification (Kandasamy *et* Kerr, 2012) for IL-6 and IL-8 cytokines secretion. The expression of many TLR-regulated genes, such as IL-6 and IL-8, were several-fold less in the LR compared to HR fibroblasts after LPS stimulation. Accordingly, levels of IL-6 and IL-8 were higher in HR compared to LR fibroblasts after LPS stimulation (Kandasamy *et* Kerr, 2012). These results suggest that the amplitude of the cells’ response to immunity stimulants may rely on a pre-existing arrangement of molecular networks that would dictate the characteristics of this response, quantitatively and possibly qualitatively.

We hypothesize that the different percentages of cells with nuclear NF-κB after LPS stimulation reflect a similar variability of fibroblasts’ response *in vivo* reflecting donor immune-status relevant information. We analyzed the expression of IL-6 and IL-8 by LPS, or poly I:C stimulated fibroblasts from donors 209 and 818 detecting secreted (supernatant), and in parallel single-cell expression by flow cytometry. Interestingly, we have observed that there were fewer cells expressing cytokines in "low responder" fibroblasts (209) than in "high responder" (818) fibroblasts for LPS stimulation. On the other hand, there was no significant difference between the responses of these same two populations of fibroblasts stimulated by poly I: C. Levels of IL-6 and IL-8 in the supernatants and populations of IL-6 and IL-8 expressing cells were consistent with the percentage of activated cells defined by the presence of NF-κB in the nucleus, suggesting the biological significance of this read-out. Finally, these high and low responses in fibroblasts do not correspond to high and low responses to LPS in the blood of the same donor (data not shown) which is consistent with previous observations (Wolf *et al.*, 2012).

Altogether, our observations suggest that fibroblasts from different donors display inter-individual variable responses to innate immune stimuli, that may translate into a stromal-specific inter-individual response variability. Indeed, stromal specific regulation of the IL-6 pathway have already been presented (Noss *et al.*, 2015; N’guyen *et al.*, 2017). Fibroblasts are a major component of the stroma where they can be exposed to microbes as well as to DAMPs in sterile inflammation and cancer. The contribution of human non-professional, tissue resident cells to protective immunity to infection were recently described (Zhang *et al.*, 2019). There is no description of fibroblasts among these non-professional cells, although the role of fibroblasts in the defense against pathogens has already been documented (Kühbacher *et al.*, 2017). Higher or lower responses to innate immune stimulants, together with a yet to be described putative qualitative variability of these responses, would probably impact the role of fibroblasts in tissue homeostasis and their dialogue with immune cells (West, 2019).

Our next steps will consider assessing the secretion level of a larger number of cytokines under these same conditions in order to better characterize the variability of this response quantitatively and qualitatively. Altogether we propose robust protocols that unravel the variability of the primary fibroblasts’ responses based on the detection of the p65 /NF-κB or imaging GFP-tagged VP26 (HSV-1). These protocols are new tools to better evaluate the functional responses of fibroblasts to a wide range of immune stimulants and their variability, a biological immune-response tissue niche whose importance has only recently come to be recognized. Indeed, subsets of fibroblasts are now directly implicated in the pathogenesis of diseases including cancer, myocardial infarction and chronic lung disease in addition to fibrotic diseases classically associated with fibroblasts.

## Author Contributions

AC, MPdM, ND, PF, TS, MH, NA, BDW: experiments and analysis, LabExMI and GvZ: resources, SLS and BDW: concept, BDW: design, data collection and writing.

## Acknowledgements

The authors thank Catherine Werts and Jong Eun Ihm for critical reading of the manuscript. BDW also thanks the members of the UTechS Photonic BioImaging (PBI) and UTechS Cytometry and Biomarkers (CB) of the Institut Pasteur for their support.

## Consortium

*Milieu Intérieur* Consortium team leaders: Laurent Abel, Matthew L. Albert, Andres Alcover, Hugues Aschard, Kalle Astrom, Philippe Bousso, Pierre Bruhns, Ana Cumano, Caroline Demangel, Ludovic Deriano, James Di Santo, Françoise Dromer, Gérard Eberl, Jost Enninga, Jacques Fellay, Magnus Fontes, Ivo Gomperts-Boneca, Milena Hasan, Serge Hercberg, Olivier Lantz, Claude Leclerc, Hugo Mouquet, Sandra Pellegrini, Stanislas Pol, Lluis Quintana-Murci, Antonio Rausell, Lars Rogge, Anavaj Sakuntabhai, Olivier Schwartz, Benno Schwikowski, Spencer Shorte, Vassili Soumelis, Frédéric Tangy, Eric Tartour, Antoine Toubert, Simon Wain-Hobson. Additional information can be found at http://www.milieuinterieur.fr/

---

Figure S1: HSV1-GFP infection of healthy and TLR3 _−/−_ fibroblasts Quantification of the percentage of infected cells after infection with 0, 01 and 0,1 MOIs. Compiled results of the mean percentages presented together for comparison: TLR3_−/−_ cells and healthy fibroblasts BJ, WI-38 and DHF. Bars plot the mean ± sd of triplicates of each experimental point.

