## Supplementary figures and images for "Unveiling inter individual variability of human fibroblast innate immune response using robust cell-based protocols"

### FIGURE S1.jpg

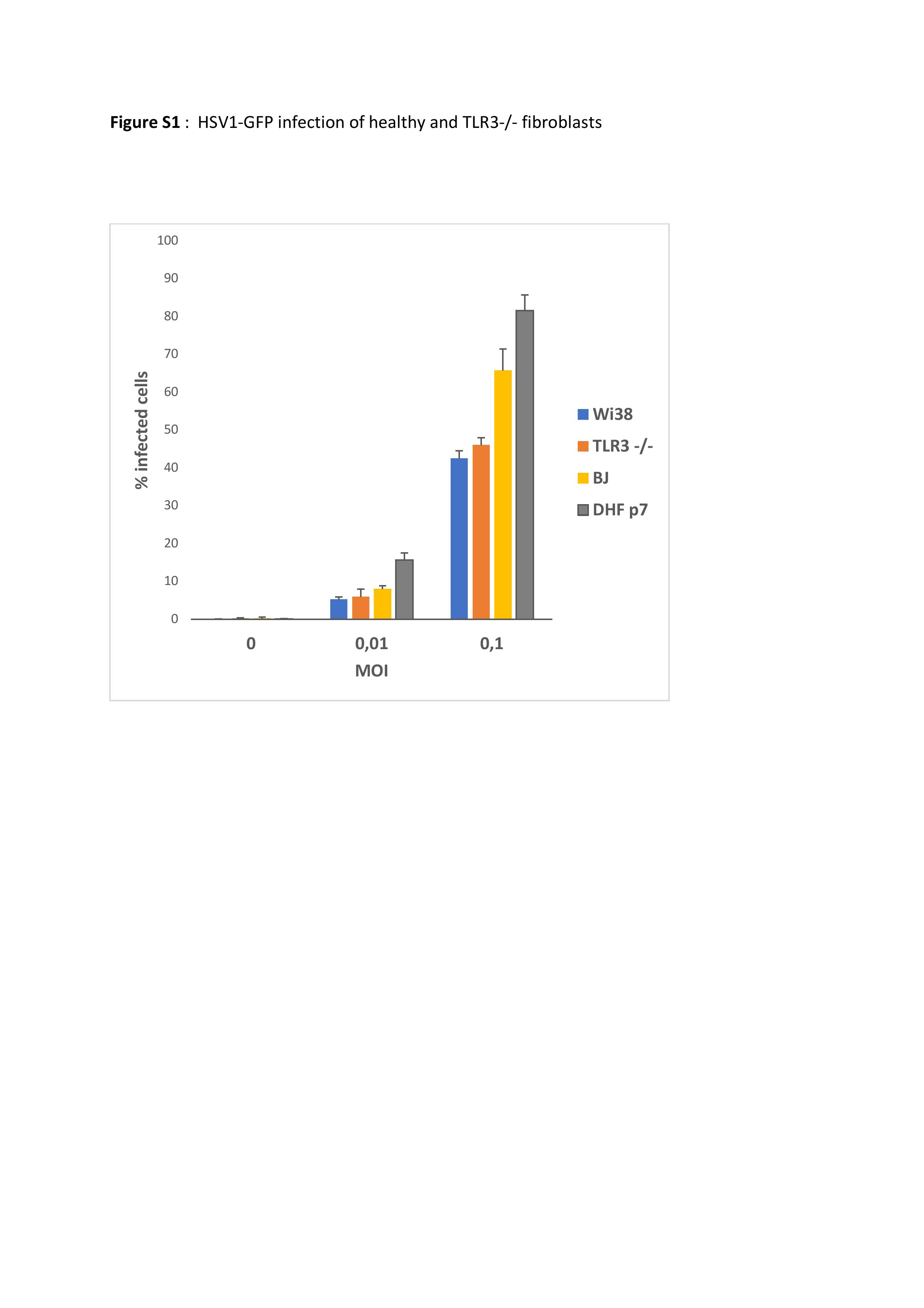
